# Identification of QTL for yield potential under heat stress conditions through a tomato MAGIC population

**DOI:** 10.1101/2025.01.23.634636

**Authors:** Ya-Ping Lin, Ehtisham Hussain, Yun-Che Hsu, Chen-Yu Lin, You-Syuan Chen, Lung-Hsin Hsu, Ching-Yuan Hung, Shu-Mei Huang, Jo-Yi Yen, Assaf Eybishitz, Peter Hanson, Seung Won Kang, Ken Hoshikawa

## Abstract

**Background:** High temperature severely limits tomato reproductive development and yield, particularly in tropical production systems. Pollen viability and fruit set are commonly used as indicators of reproductive heat tolerance, but much of the supporting evidence has been obtained under controlled conditions. Their relevance to breeding under variable open-field environments remains less clear. This study investigated reproductive performance and its genetic basis under open-field heat stress using a multi-parent advanced generation inter-cross (MAGIC) population.

**Results:** We developed a MAGIC population from eight *Solanum lycopersicum* parents combining heat tolerance and multiple disease resistances and evaluated 262 F_5:6_ lines under natural open-field heat stress. Flower number showed the highest heritability (0.64), followed by pollen viability (0.56), fruit number (0.48), and days to flowering (0.25), whereas fruit set showed negligible heritability (<0.01). Despite its moderate heritability, pollen viability was only weakly correlated with fruit set (*r* = 0.22). In contrast, fruit number was strongly correlated with fruit set (*r* = 0.86), supporting fruit number as a practical selection criterion for reproductive performance under open-field heat stress. Restriction enzyme-associated DNA sequencing generated 16,350 high-quality SNPs for genome-wide association analysis, which identified multiple QTLs for reproductive traits, several overlapping previously reported loci. A region on chromosome 6 at approximately 31–33 Mb was particularly notable because it was associated with multiple reproductive traits and coincided with a previously reported heat-responsive QTL region. Candidate-gene analysis highlighted floral developmental regulation, represented by positional candidates *SlAGO7*, *ENO*, and *SlWUS*, and cellular and protein homeostasis, supported by direct candidates including MON1, ECM29, and ubiquitin-related genes.

**Conclusions:** Selection criteria established under controlled environments may not fully predict reproductive performance under open-field heat stress. Fruit number provides an integrative and moderately heritable indicator of realized reproductive success under field conditions. The identified QTLs and candidate genes provide targets for validation and development of heat-tolerant tomatoes adapted to tropical open-field production.

## Background

Tomato (*Solanum lycopersicum*) is a globally important crop for both fresh consumption and processing. Its production is increasingly threatened by heat stress, which is intensifying due to climate change [1]. Optimal growth occurs at 26–28°C (day) and 22°C (night), while temperatures above 32°C (day) or 26°C (night) disrupt plant development [2]. Heat stress impairs photosynthesis by reducing the maximum quantum yield (Fv/Fm) of photosystem II, damaging chlorophyll function and overall capacity [3, 4]. It destabilizes membranes, proteins, and enzymes, and induces oxidative stress through reactive oxygen species (ROS) accumulation [5]. The reproductive phase is especially sensitive, with high temperatures affecting microsporogenesis, pollen viability, and fruit set [6–11]. Prolonged exposure to mild heat (32°C, 70% RH, 14-h light) can reduce pollen viability by 86%, pollen number by 56%, and female fertility by 39% [12]. The rising frequency of heatwaves in major production regions exacerbates these impacts, highlighting the urgency of identifying the genetic basis of heat tolerance for breeding resilient cultivars [13].

Domestication has narrowed genetic diversity in cultivated tomatoes, limiting stress tolerance [14–17]. Breeding programs have historically focused on yield and fruit quality under optimal conditions, often at the expense of resilience [18]. Wild relatives such as *Solanum pimpinellifolium* retain key adaptive traits, including high pollen viability, sustained chlorophyll content, and stable photochemistry under heat stress [4, 19, 20], making them valuable for breeding. However, introgression is hindered by linkage drag and the need to restore desirable fruit traits. The polygenic nature of heat tolerance, involving flowering time, pollen viability, and fruit set, further complicates breeding, necessitating quantitative trait locus (QTL) mapping and targeted trait selection.

In the past decade, QTL mapping has identified genomic regions underlying heat tolerance. Previously research reported *qPN7* and *qPV11* on chromosomes 7 and 11 [8], explaining 18.6% and 36.3% of variation in pollen number and viability. QTL *AB7.1_T2* on chromosome 7 for pollen viability and several loci on chromosome 2 for flower number, fruit number, and fruit set were identified [20]. In addition, a QTL hub on chromosome 3 (53–65 Mb) controlling multiple reproductive traits was also found [21]. Meanwhile, a locus on chromosome 1 (82–86 Mb) linked to relative electrical conductivity and chlorophyll content was detected [4]. These studies highlight chromosomes 1, 2, and 3 as key regions for heat tolerance.

However, most mapping has relied on bi-parental populations, limiting allelic diversity and resolution. Multi-parent advanced generation intercross (MAGIC) populations overcome these constraints by incorporating multiple founders, increasing recombination, and balancing allele frequencies. The first tomato MAGIC population, developed by Pascual et al. (2015) from eight *S. lycopersicum* accessions, increased recombination frequency by 87% compared to bi-parental designs, enabling fine mapping of fruit weight QTLs [22]. This resource has also been used to study genotype × environment (G × E) interactions, aiding climate-resilient breeding [23]. The ToMAGIC population, developed from *S. lycopersicum* var. *cerasiforme* and *S. pimpinellifolium*, captured broader genetic diversity for discovering loci associated with fruit size, pigmentation, and leaf morphology [24]. By integrating diverse founder genomes, MAGIC populations reduce candidate polymorphisms and improve mapping precision, making them powerful tools for dissecting complex traits like heat tolerance.

In this study, aligned with the World Vegetable Center’s breeding objectives, heat tolerance is defined as the ability to maintain high reproductive performance, i.e. yield potential, under heat stress. A MAGIC population derived from heat-tolerant crossing disease-resistant *S. lycopersicum* lines was evaluated and subjected to genome-wide association studies (GWAS) to identify loci linked to reproductive traits. These results identify key QTLs associated with maintaining yield potential under heat stress, providing valuable targets for breeding heat-resilient cultivars adapted to climate change.

## Materials and Methods

### Development of the MAGIC population

The MAGIC population was designed with two primary objectives: (1) to identify genes underlying heat tolerance in tomato, and (2) to develop elite heat-tolerant lines for breeding new varieties suited to tropical and subtropical regions. Parent selection also considered traits beyond heat tolerance. In tropical production systems, begomoviruses causing tomato yellow leaf curl disease (TYLCD) are a major constraint, and high temperatures often coincide with high TYLCD pressure, complicating field evaluation of heat-related traits. Therefore, candidate founder parents were screened with molecular markers for the *Ty-1/3* gene, which confers tolerance to a broad spectrum of begomoviruses. Parents were genotyped for other disease resistance genes including *Bwr-*12 (tolerance to bacterial wilt disease caused by *Ralstonia solanacearum,* a common soilborne pathogen in the tropics) and *Ph-*3 (a gene conferring resistance to some strains of *Phytophthora infestans* causing late blight). The marker information is listed in Table S1. Candidate parents were also evaluated for horticultural (vigor, plant habit, maturity) and fruit (weight, color, cracking) traits.

### Evaluation and selection of the MAGIC parental lines

Ten breeding materials (eight *S. lycopersicum* and two *S. pimpinellifolium*) were evaluated for pollen viability and high temperature fruit set in a greenhouse from August to October 2017 at the World Vegetable Center, Shanhua, Tainan. Mean day and night temperatures during this period were 35.4℃, and 28.1℃, respectively; the highest temperature was 43.4℃ and the lowest was 24.4℃. The candidate heat-tolerant parents were selected from literature review and breeders’ experience (Table 1). We used a randomized complete block design with three blocks and four plants per plot. Pollen was collected from the most recently opened flowers on the second to sixth inflorescences of the main stem for each plant. The pollen viability was assessed using a protocol adapted from Abdul-Baki [25]. Briefly, anther cones were removed from the flowers, opened, and squashed using tweezers. The released pollen was then stained by vortexing in 150 µL of 0.5% acetocarmine solution. A 10 µL aliquot of the stained pollen was placed on a glass slide, covered with coverslip, and observed under 40x magnification. Viable pollen grains (stained red) and non-viable pollen grains (transparent) were counted. Fruit set was determined as the ratio of the total number of fruits to the total number of flowers on the second to sixth inflorescences of the main stem.

**Table 1.**
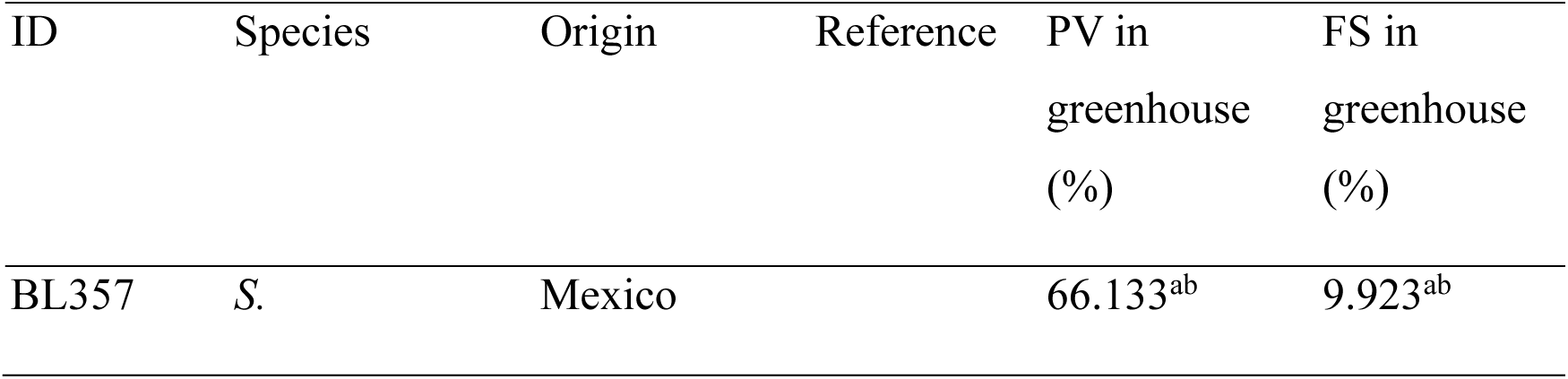
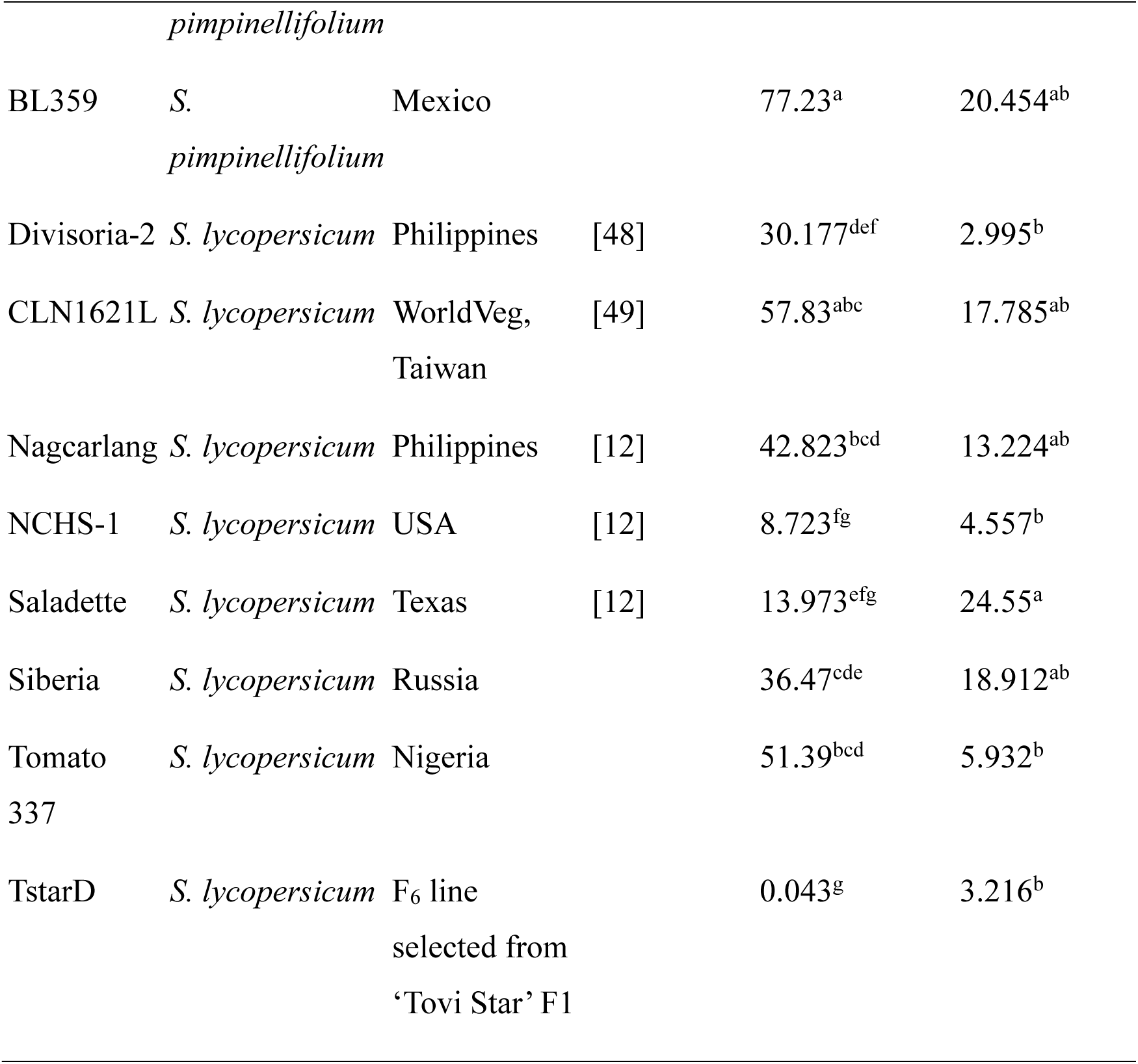
Candidate parental lines evaluated for pollen viability and fruit set in the greenhouse.

A detailed workflow of the crossing process is presented in Fig. 1A. Eight cultivated tomato lines were selected as the foundation parents of the MAGIC population based on the greenhouse trial. To develop F_1_ hybrids, we initially crossed heat-tolerant lines with disease-resistant lines. To enhance recombinant events, crossing the double crosses was performed by growing 120 plants each of CLN4430 and CLN4433. Each plant in both populations was used once as a male and once as a female in crosses. This process resulted in a total of 240 intercrosses between the double hybrids, allowing for the mixing of all eight genomes. The progeny derived from these crosses was advanced through successive generations using the single-seed descent method. To mitigate the risk of losing progeny under open-field conditions, four plants per progeny were initially planted. To preserve genetic integrity and avoid the effects of artificial selection on gene flow and pool structure, the first plant from each progeny was retained and harvested for the next generation. Additionally, superior plants were identified, selected, and harvested. As a result, 262 progenies were ultimately maintained and advanced to the F_5:6_ generation.

**Fig. 1.**
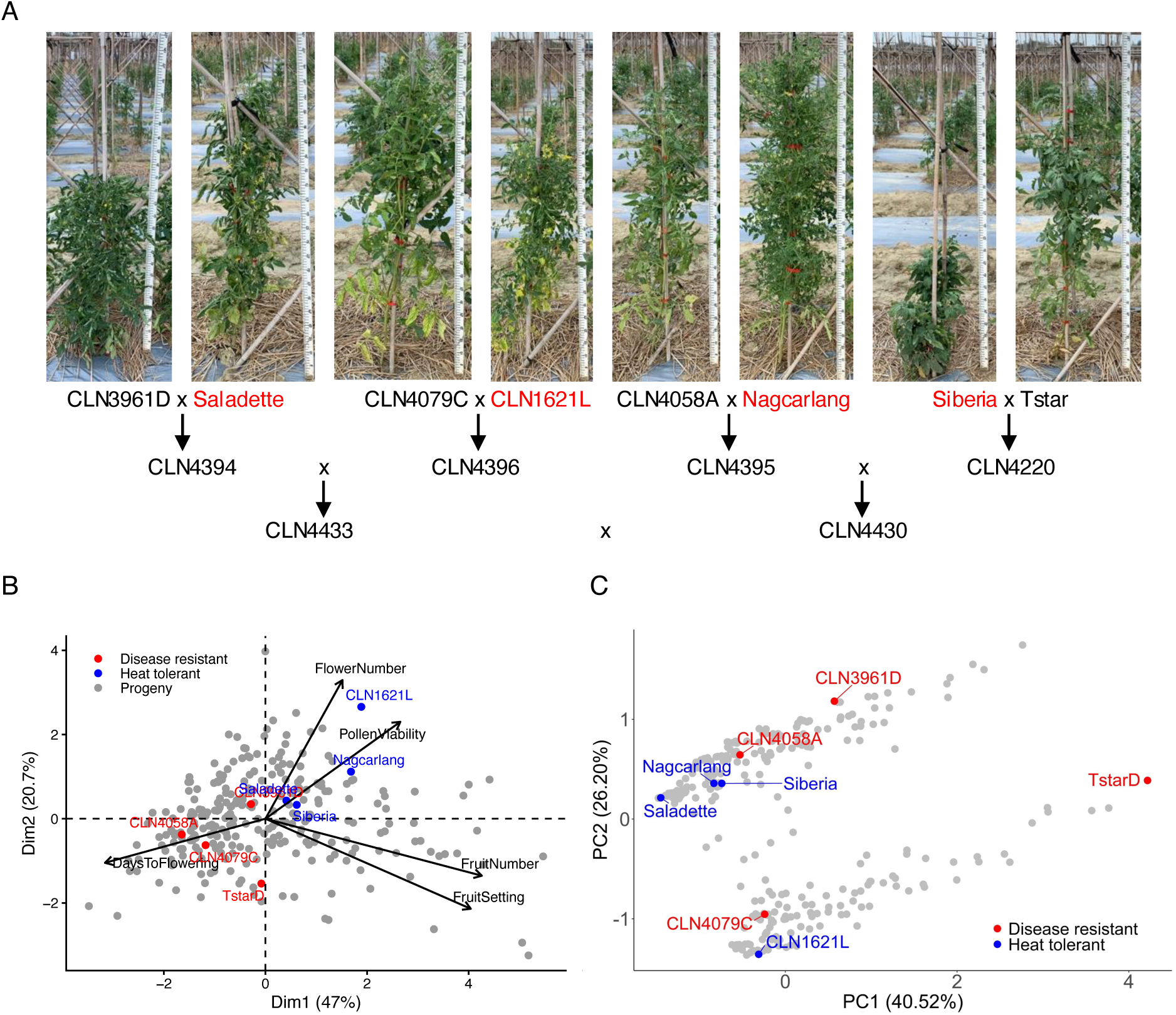
Overview of the tomato MAGIC population. **A** Eight parental lines used to develop the MAGIC population and the crossing scheme. **B** Principal component analysis (PCA) of five reproductive traits. **C** Principal coordinate analysis (PCoA) showing genetic relationships among the parental lines and population structure within the MAGIC population. In **B** and **C**, red, blue, and grey points represent disease-resistant parents, heat-tolerant parents, and MAGIC progeny, respectively.

### Phenotyping of the MAGIC population

For phenotypic evaluation, a total of 262 F_5:6_ lines were planted in two replications. Four plants per progeny per replication were evaluated in the field under natural heat stress conditions during the summer season in 2021 at the World Vegetable Center, Headquarters, Shanhua, Tainan. The seeds were sowed on the 15^th^ of July; the seedlings were transplanted on the 12^th^ of August; fruit was harvested on the 18^th^ of October. The daily mean, maximum, and minimum temperatures were recorded at 27.9℃, 37.2℃, and 19.7℃, respectively. We evaluated lines for multiple traits, including days to flowering (DF), pollen viability (PV), flower number (FLN), fruit number (FRN), and fruit set (FS). Days to flowering were recorded as the days until 50% of the plants had flowered. Flower number, fruit number, and fruit set were counted and estimated on the second to sixth inflorescences of the main stem.

Pollen viability was measured using Ampha Z32 Impedance Flow Cytometer (Amphasys AG, Switzerland) to determine the number of viable pollen grains and total pollen count. Pollen viability tests were conducted on the second and fourth inflorescences, with two open flowers randomly collected from two plants within each plot. The anther cones were removed from the flowers, placed in a 2.0 mL Eppendorf tube containing 750 µL of AmphaFluid 6 (AF6) buffer (Amphasys AG, Switzerland), and vigorously scraped along the tube bottom to release pollen until the solution became turbid. The turbid solution was filtered, and an additional 750 µL of AF6 buffer was used to wash any remaining pollen on the filter membrane. After the measurement, pollen viability was quantified using AmphaSoft 2.1.6 software (Amphasys AG, Switzerland).

The correlation and principal component analysis (PCA) among traits were conducted with the R cor() and prcomp() functions, respectively, and visualized by corrplot and ggplot2 packages, respectively [26–28].

### Genotyping

For the eight parents, young leaves of individual seedlings were collected for 10x coverage of the whole genome resequencing. The sequencing was outsourced using Illumina HiSeq 2500 platform of paired-end reads. After data was retrieved, we processed the raw reads following the GATK Best Practice to perform the reference genome-based single nucleotide polymorphism (SNP) calling [29]. In brief, we used SolexaQA++ “dynamictrim” to clean the reads [30]. The cleaned reads were mapped to SL4.0 using BWA [31, 32]. The individual SNPs were called by gatk functions “HaplotypeCaller” and “GenotypeGVCFs” and then merged by Picard MergeVcfs [33]. The SNP variants were filtered with the threshold of sequencing quality > 20 and mapping quality > 40. Then, we removed all the missing and heterozygous SNPs.

Meanwhile, young leaves of the F_5:6_ progenies were collected, and we divided the samples evenly into three groups to prepare the *Ape*KI-digested DNA libraries. The sequencing was outsourced using Illumina HiSeq 2500 platform of paired-end reads. After data retrieval, we processed the raw reads following the similar procedure with what we did for the whole genome sequencing. In brief, we used Stacks function “process_radtags” to demultiplex the data and to check quality scores of the reads [34]. The SNPs were developed following the GATK Best Practice and filtered with the threshold of sequencing quality > 20 and mapping quality > 40 [29]. Samples with more than 50% missing sites and SNPs with more than 25% missing sites were removed from the dataset. We aligned the progeny SNP matrix with the parents SNP matrix, and the missing sites were imputed using Beagle 5.5 [35].

### Mapping of the reproductive traits

Phenotypic distributions were examined for the five reproductive traits using histograms and kernel density estimates. The normal distribution expected from the observed mean and standard deviation was overlaid for visual comparison, and skewness was calculated to assess departures from symmetry using skewness_fun() function in R. To reduce skewness and the influence of extreme observations, phenotypic values were log10-transformed prior to GWAS analysis. For traits containing zero values, log10(x + 1) transformation was applied to retain zero-valued observations.

The genetic distance of the parental lines and the progeny was first estimated in TASSEL v5.2.89 [36]. Principal coordinate analysis (PCoA) was performed using the prcomp() function in R, with the genetic distance matrix obtained from TASSEL as input. The result was visualized using R ggplot2 package [27]. Since this population was generated from multiple parents, we considered it germplasm and conducted association mapping using the Genome-wide Efficient Mixed Model Association (GEMMA) software [37]. GEMMA utilizes a linear mixed model framework that effectively controls for confounding due to both population stratification and sample relatedness. In this study, we used a linear mixed model treating SNPs as fixed and the kinship matrix estimated by GEMMA as a random effect. The heritability was estimated using the Bayesian Sparse Linear Mixed Model implemented in GEMMA software [38]. The candidate SNPs were defined as *P score* < 0.001 due to the limitation of the replication in the field evaluation. FDR-adjusted *P* values were calculated using the Benjamini–Hochberg procedure in R and reported as complementary measures of statistical evidence rather than as the primary threshold for declaring significant marker–trait associations.

To define QTLs and facilitate the development of practical markers, significant SNPs located within 5 Mb of each other were grouped into single QTL regions. This 5 Mb window was determined based on the uneven and often extensive linkage disequilibrium (LD) decay observed in tomato populations [39]. From each QTL group, the SNP with the most significant association was chosen first as the representative marker for downstream analyses. A direct candidate gene was defined when a significantly associated SNP was located directly within the gene. The QTLs of reproductive traits in the four previous studies [8, 11, 20, 21] were browsed on the SL4.0 genome using Genome Browser hosted on the Sol Genomics Network (https://solgenomics.sgn.cornell.edu/) to confirm the consistency with the QTL identified in this study.

## Results

### Evaluation of candidate MAGIC parental lines

To assess heat tolerance, ten tomato materials were evaluated for pollen viability and fruit set, revealing significant genotypic effects for both traits (*P value* < 0.01). Among the *S. lycopersicum* accessions, the highest to lowest pollen viability was observed in the following order: CLN1621L, Tomato 337, Nagcarlang, Siberia, Divisoria-2, Saladette, NCHS-1, and TstarD (Table 1). However, the rankings for fruit set differed, with Saladette, Siberia, CLN1621L, and Nagcarlang showing superior performance. As expected, the two small-fruited *S. pimpinellifolium* accessions exhibited higher pollen viability and fruit set than the cultivated *S. lycopersicum*. Given the goal of developing high-yielding tomato varieties without the linkage drag associated with wild tomatoes, fruit set among *S. lycopersicum* was prioritized as the key selection criterion for identifying heat-tolerant parental lines. Based on this criterion, Saladette, Siberia, CLN1621L, and Nagcarlang emerged as the most promising heat tolerant candidate although each is begomovirus and bacterial wilt susceptible. Consequently, four additional multiple disease resistant lines with superior fruit and horticultural traits were also included as parents.

Field trials further elucidated the heat tolerance of the parental lines under open-field conditions that reflected realistic agricultural practices. CLN1621L and Nagcarlang exhibited superior pollen viability in open-field environments (Table 2). Additionally, Nagcarlang and Saladette demonstrated the highest fruit set and fruit number among the materials tested (Table 2), underscoring Nagcarlang’s robustness across multiple experiments essential for reproductive success under heat stress. Interestingly, TstarD exhibited high fruit set and fruit number in field conditions despite its low pollen viability. This contrasted with its greenhouse performance, where it ranked among the lowest for both fruit set and pollen viability, indicating that TstarD is highly sensitive to environmental conditions. These findings highlight the importance of considering environmental adaptability when selecting heat-tolerant parental lines for breeding programs.

**Table 2.**
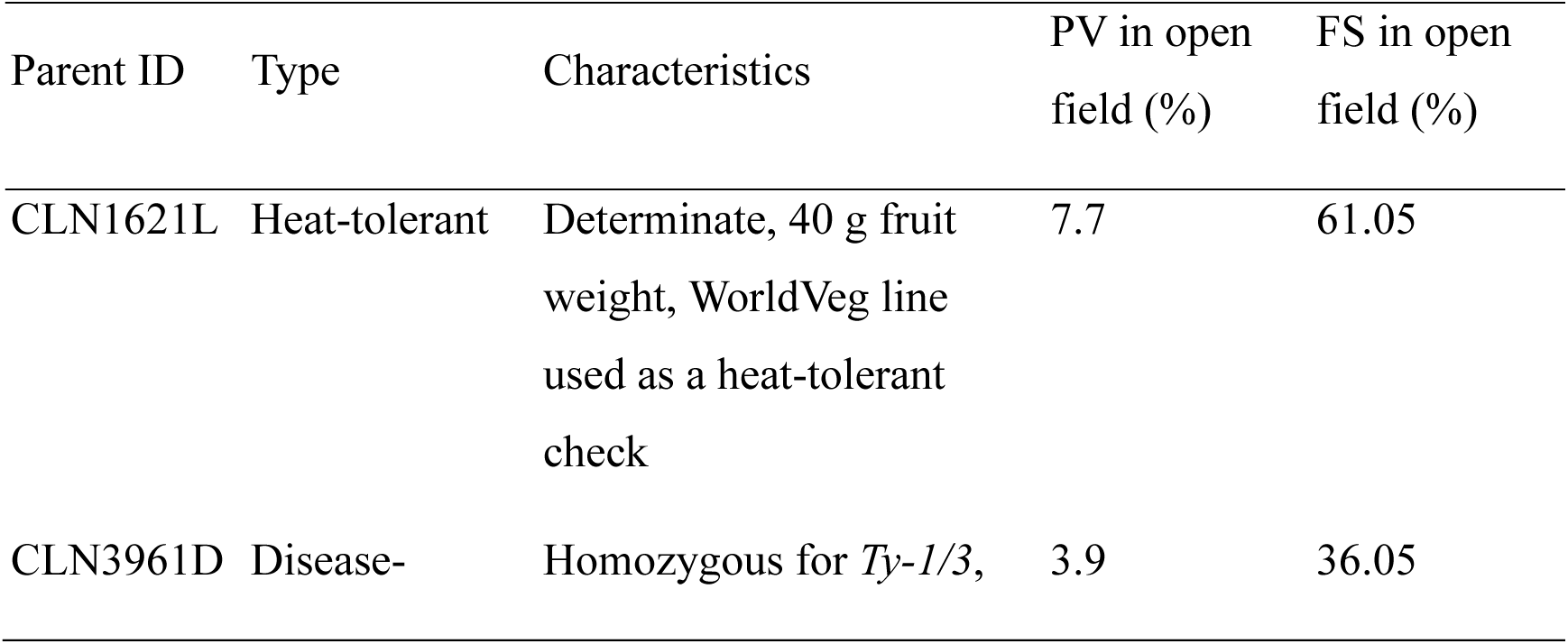
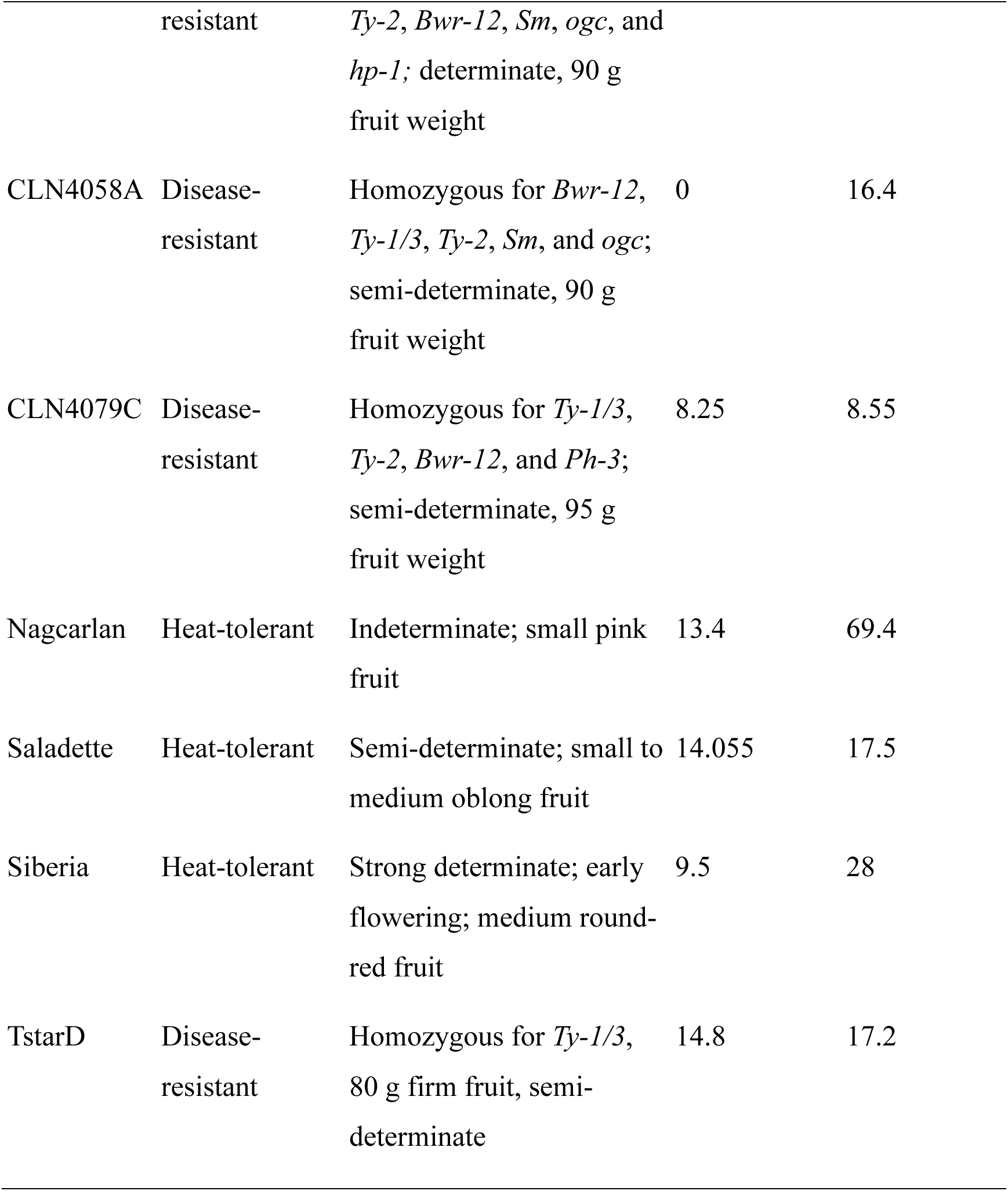
Horticultural characteristics, marker information, and phenotypes of the MAGIC parental lines.

### Fruit number as a selection index for heat tolerance under open-field conditions

Days to flowering ranged from 10.5 to 44.0 days, pollen viability from 1.35 to 69.40%, flower number from 1.5 to 12.1, fruit number from 0 to 45.75, and fruit set from 0 to 56.95% (Fig. 2). Analysis of the relationships among reproductive traits under heat stress revealed that days to flowering was negatively correlated with all other reproductive traits, with Pearson’s correlation coefficients (*r*) ranging from −0.14 to −0.38 (Fig. S1), indicating that earlier flowering was generally associated with better reproductive performance under high-temperature conditions. Among the reproductive traits, fruit set was most strongly correlated with fruit number (*r* = 0.86, *P* < 0.001), followed by a moderate negative correlation with days to flowering (*r* = −0.36, *P* < 0.001). In contrast, pollen viability showed only a weak positive correlation with fruit set (*r* = 0.22, *P* < 0.001).

**Fig. 2.**
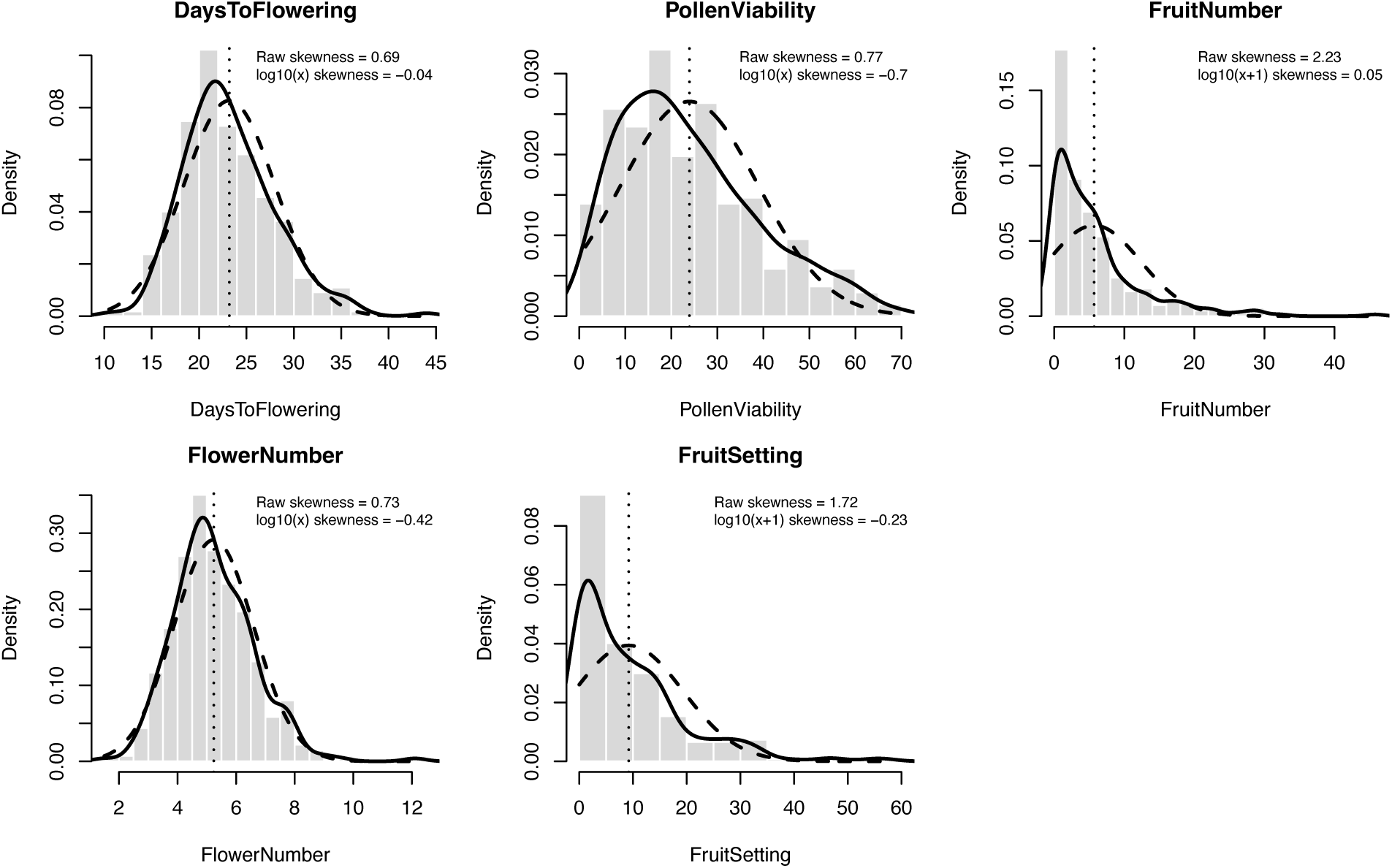
Phenotypic distributions of reproductive traits in the tomato MAGIC population under open-field heat-stress conditions. Histograms represent the original phenotypic values for five reproductive traits. Skewness before and after log10 transformation is shown for each trait to assess changes in distributional symmetry.

Consistent with these pairwise relationships, PCA revealed similar loading directions for fruit number and fruit set, whereas pollen viability and flower number were oriented in a different direction, indicating partially distinct patterns of variation among these reproductive traits. Days to flowering showed an opposing loading direction to fruit number and fruit set. The first two principal components explained 67.7% of the total phenotypic variation, with PC1 and PC2 accounting for 47.0% and 20.7%, respectively (Fig. 1B). Together, the correlation and multivariate analyses indicate that fruit number was more closely associated with realized fruit set than pollen viability under open-field heat stress, supporting its potential as a practical selection index for reproductive heat tolerance.

### Genetic diversity and population structure of this population

Whole-genome resequencing of the eight parental lines identified 4,075,083 SNPs, providing a high-density genomic reference for the MAGIC population. Following SNP filtering and genotype imputation, 16,350 high-quality SNPs were retained for subsequent GWAS (Fig. 3). The average marker interval across the genome was approximately 48 kb, ranging from 17 kb on chromosome 6 to 195 kb on chromosome 1 (Table 3). Principal coordinate analysis (PCoA) revealed relatively close genetic relationships among most heat-tolerant parents, with CLN1621L showing greater differentiation, whereas the disease-resistant parents exhibited broader genetic diversity (Fig. 1C). Within the MAGIC population, PCoA revealed two partially differentiated groups with substantial admixture, indicating residual population structure following intercrossing and recombination during population development (Fig. 1C). This structure was accounted for in the subsequent GWAS using a kinship matrix.

**Fig. 3.**
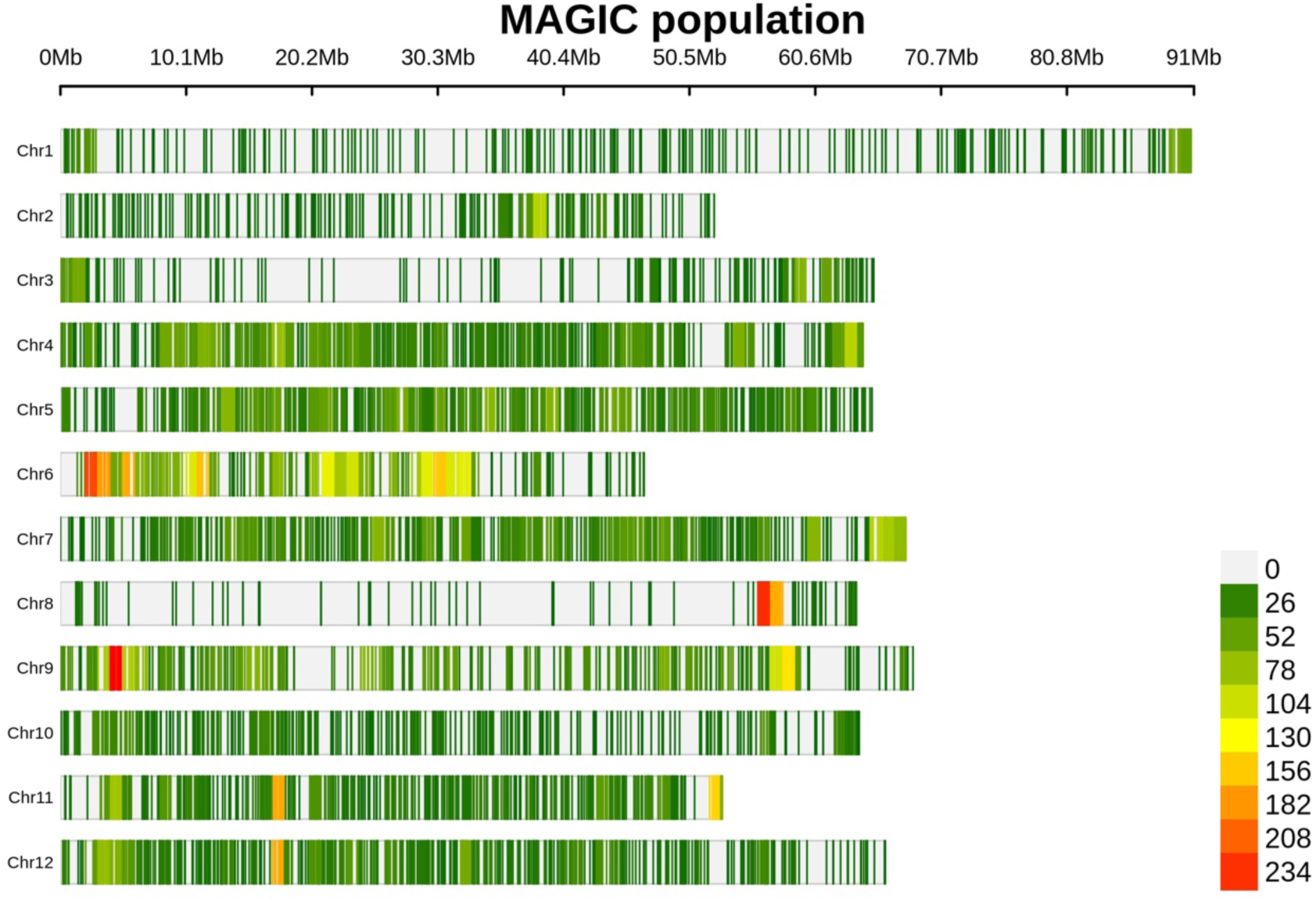
Genome-wide distribution of SNP markers in the tomato MAGIC population. SNP density across the 12 tomato chromosomes was calculated in 1-Mb windows. Colors indicate the number of SNPs per window, ranging from low (green) to high (yellow–red) density; grey regions indicate windows without SNP markers. Chromosome positions are shown in megabases (Mb).

**Table 3.**
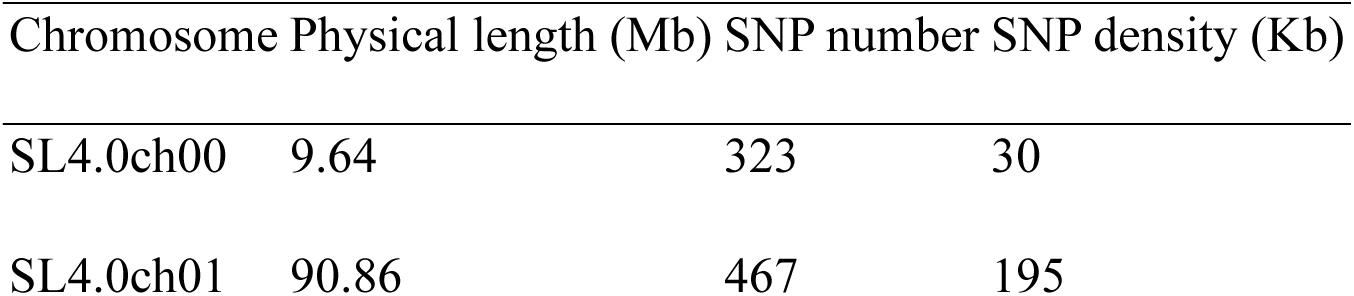
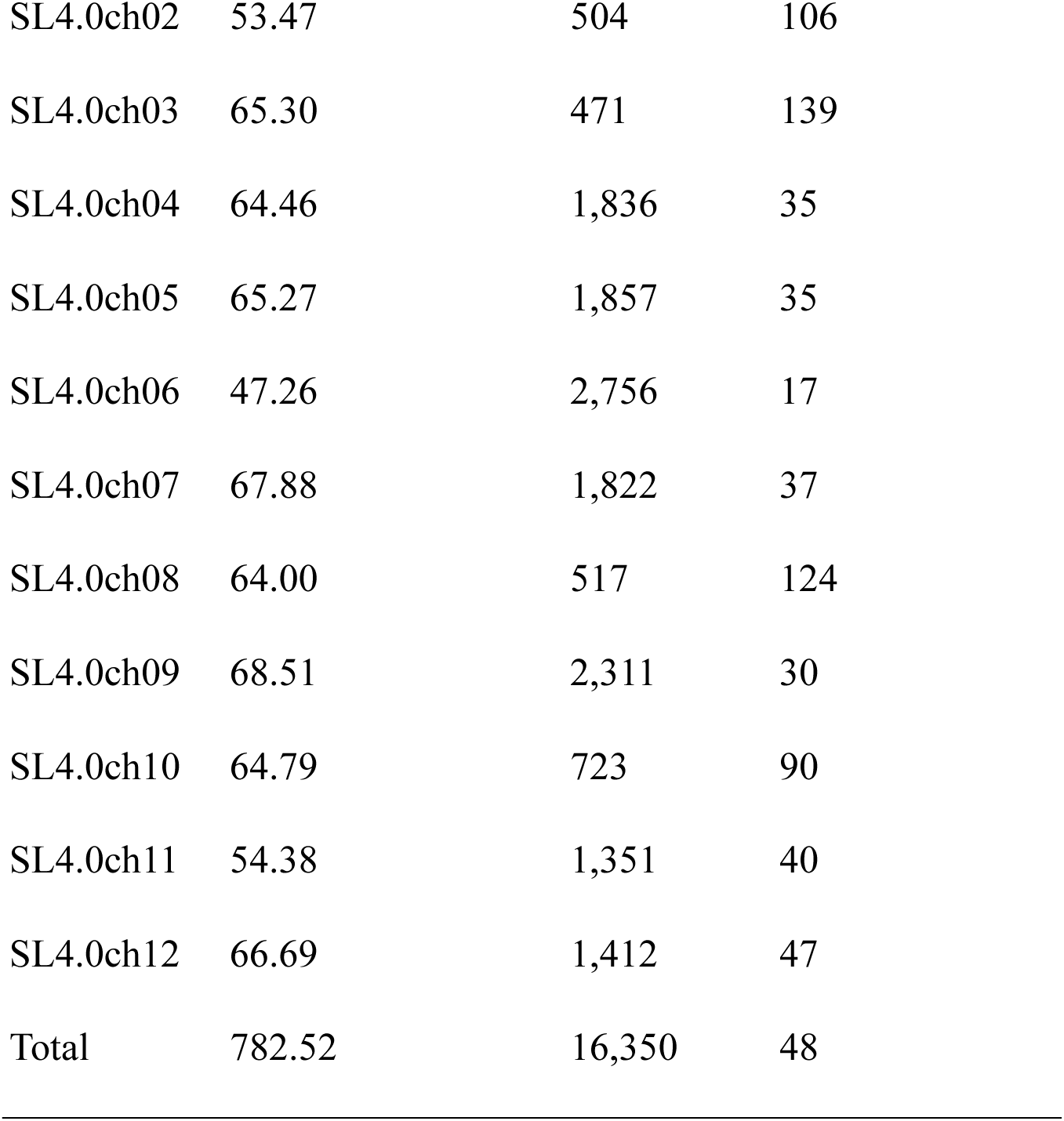
SNP density across the genome.

### QTLs for reproductive traits under heat stress

Given the genetic diversity among the parental lines, the MAGIC population was treated as a diverse association panel for GWAS. In total, 62 significant trait–SNP associations, representing 54 unique SNPs, were detected for five reproductive traits under heat stress conditions (Fig. 4; Fig. S2; Table S3). Of these associations, 20 were detected for days to flowering, 8 for pollen viability, 18 for flower number, 12 for fruit number, and 4 for fruit set. Three SNPs were associated with multiple reproductive traits: SL4.0CH01_7511997 with days to flowering, pollen viability, and flower number; SL4.0CH01_24105448 with pollen viability, flower number, and fruit number; and SL4.0CH01_62158112 with days to flowering, flower number, and fruit number. In addition, SL4.0CH01_63692426 was shared between days to flowering and fruit number, whereas SL4.0CH04_61821975 was associated with both fruit number and fruit set.

**Fig. 4.**
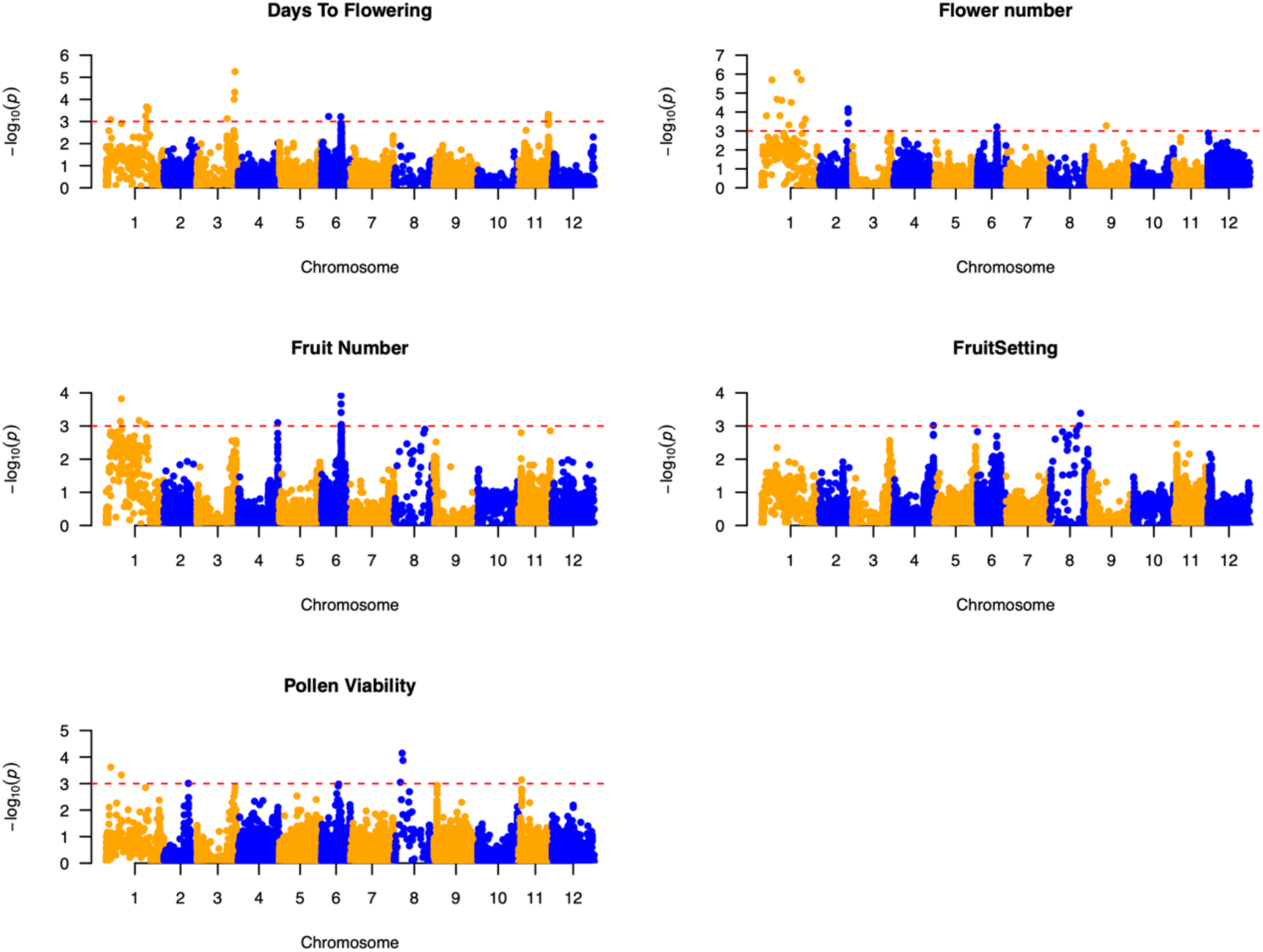
Manhattan plots of genome-wide association studies for five reproductive traits in the tomato MAGIC population under open-field heat-stress conditions. Manhattan plots show marker–trait associations for days to flowering, pollen viability, flower number, fruit number, and fruit set across the 12 tomato chromosomes. The horizontal dashed line indicates the significance threshold used to identify significant SNPs.

To account for the extensive heterochromatic regions and large linkage intervals in tomato, significant SNPs for each trait located within 5 Mb of one another were grouped into QTL regions. This resulted in 29 trait-specific QTLs, including 7 for days to flowering, 5 for pollen viability, 9 for flower number, 5 for fruit number, and 3 for fruit set (Table 4). Several genomic regions showed associations with multiple reproductive traits. Notably, regions on chromosome 1 contained overlapping QTLs for days to flowering, pollen viability, flower number, and fruit number. A region on chromosome 6 at approximately 31.6–32.5 Mb was also associated with days to flowering, flower number, and fruit number, suggesting that these genomic regions may contribute to the coordinated genetic regulation of reproductive responses under heat stress.

**Table 4.**
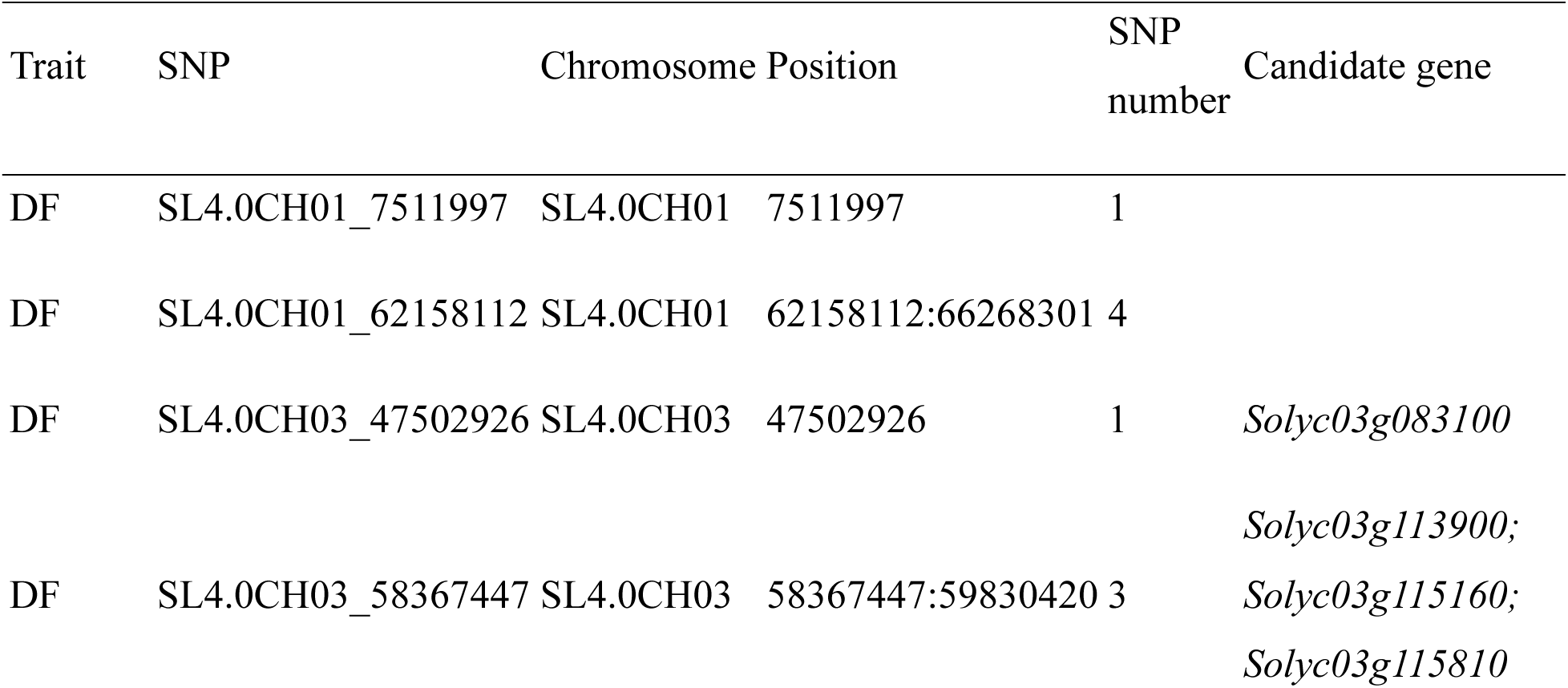
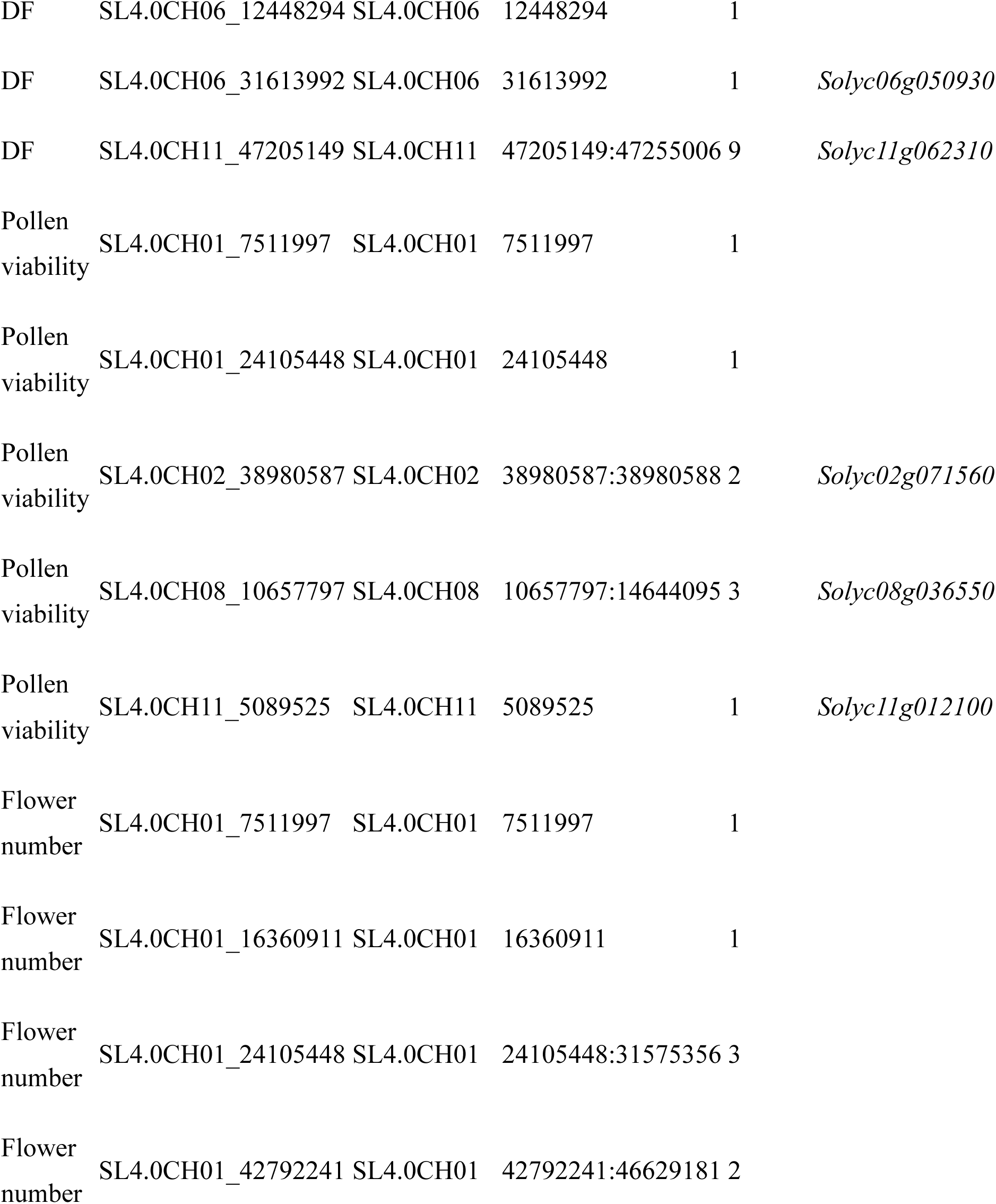
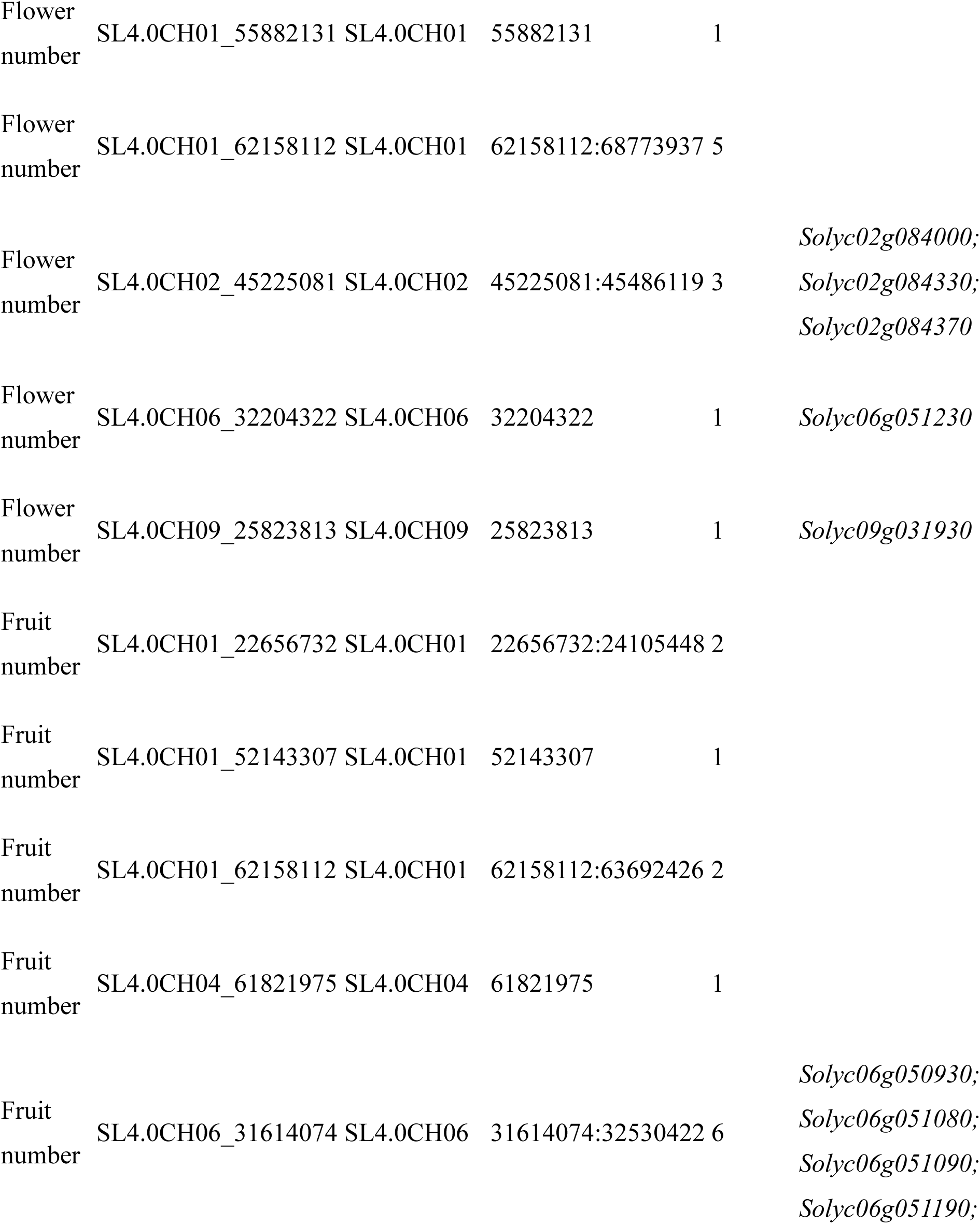
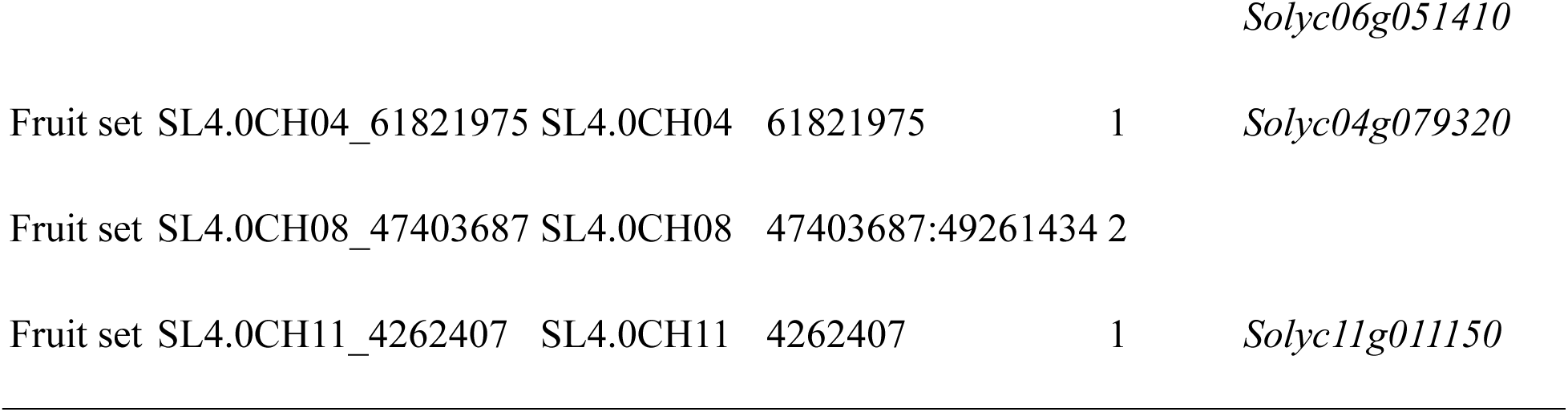
QTL regions associated with reproductive traits under heat stress, defined by grouping significant SNPs within 5 Mb in the tomato MAGIC population.

| Trait | SNP | Chromosome | Position | SNP number | Candidate gene |
| --- | --- | --- | --- | --- | --- |
| DF | SL4.0CH01_7511997 | SL4.0CH01 | 7511997 | 1 |  |
| DF | SL4.0CH01_62158112 | SL4.0CH01 | 62158112:66268301 | 4 |  |
| DF | SL4.0CH03_47502926 | SL4.0CH03 | 47502926 | 1 | *Solyc03g083100* |
| DF | SL4.0CH03_58367447 | SL4.0CH03 | 58367447:59830420 | 3 | *Solyc03g113900;*<br>*Solyc03g115160;*<br>*Solyc03g115810* |

| Trait | SNP | Chromosome | Position | SNP<br>number | Candidate gene |
| --- | --- | --- | --- | --- | --- |
| DF | SL4.0CH06_12448294 | SL4.0CH06 | 12448294 | 1 |  |
| DF | SL4.0CH06_31613992 | SL4.0CH06 | 31613992 | 1 | *Solyc06g050930* |
| DF | SL4.0CH11_47205149 | SL4.0CH11 | 47205149:47255006 | 9 | *Solyc11g062310* |
| Pollen<br>viability | SL4.0CH01_7511997 | SL4.0CH01 | 7511997 | 1 |  |
| Pollen<br>viability | SL4.0CH01_24105448 | SL4.0CH01 | 24105448 | 1 |  |
| Pollen<br>viability | SL4.0CH02_38980587 | SL4.0CH02 | 38980587:38980588 | 2 | *Solyc02g071560* |
| Pollen<br>viability | SL4.0CH08_10657797 | SL4.0CH08 | 10657797:14644095 | 3 | *Solyc08g036550* |
| Pollen<br>viability | SL4.0CH11_5089525 | SL4.0CH11 | 5089525 | 1 | *Solyc11g012100* |
| Flower<br>number | SL4.0CH01_7511997 | SL4.0CH01 | 7511997 | 1 |  |
| Flower<br>number | SL4.0CH01_16360911 | SL4.0CH01 | 16360911 | 1 |  |
| Flower<br>number | SL4.0CH01_24105448 | SL4.0CH01 | 24105448:31575356 | 3 |  |
| Flower<br>number | SL4.0CH01_42792241 | SL4.0CH01 | 42792241:46629181 | 2 |  |
| Flower<br>number | SL4.0CH01_55882131 | SL4.0CH01 | 55882131 | 1 |  |
| Flower<br>number | SL4.0CH01_62158112 | SL4.0CH01 | 62158112:68773937 | 5 |  |
| Flower<br>number | SL4.0CH02_45225081 | SL4.0CH02 | 45225081:45486119 | 3 | *Solyc02g084000;*<br>*Solyc02g084330;*<br>*Solyc02g084370* |
| Flower<br>number | SL4.0CH06_32204322 | SL4.0CH06 | 32204322 | 1 | *Solyc06g051230* |
| Flower<br>number | SL4.0CH09_25823813 | SL4.0CH09 | 25823813 | 1 | *Solyc09g031930* |
| Fruit<br>number | SL4.0CH01_22656732 | SL4.0CH01 | 22656732:24105448 | 2 |  |
| Fruit<br>number | SL4.0CH01_52143307 | SL4.0CH01 | 52143307 | 1 |  |
| Fruit<br>number | SL4.0CH01_62158112 | SL4.0CH01 | 62158112:63692426 | 2 |  |
| Fruit<br>number | SL4.0CH04_61821975 | SL4.0CH04 | 61821975 | 1 |  |
| Fruit<br>number | SL4.0CH06_31614074 | SL4.0CH06 | 31614074:32530422 | 6 | *Solyc06g050930;*<br>*Solyc06g051080;*<br>*Solyc06g051090;*<br>*Solyc06g051190;* |
|  |  |  |  |  | *Solyc06g051410* |
| Fruit set | SL4.0CH04_61821975 | SL4.0CH04 | 61821975 | 1 | *Solyc04g079320* |
| Fruit set | SL4.0CH08_47403687 | SL4.0CH08 | 47403687:49261434 | 2 |  |
| Fruit set | SL4.0CH11_4262407 | SL4.0CH11 | 4262407 | 1 | *Solyc11g011150* |

The heritability of the five reproductive traits varied considerably, ranging from < 0.01 to 0.64, indicating substantial differences in the relative contributions of genetic and environmental effects. Flower number showed the highest heritability (0.64), followed by pollen viability (0.56) and fruit number (0.48), suggesting moderate to strong genetic control of these traits under heat stress. Days to flowering showed relatively low heritability (0.25), whereas fruit set had negligible heritability (< 0.01), indicating that variation in fruit set was predominantly influenced by environmental effects. Although pollen viability was moderately heritable, its relatively weak correlation with fruit set suggests that it may not adequately predict reproductive success under field heat stress. In contrast, the combination of moderate heritability and a strong association with fruit set supports fruit number as a useful selection criterion for improving reproductive performance under heat stress. Flower number, with the highest heritability, may provide an additional genetically stable trait for selection, although its relationship with final reproductive success should also be considered.

### Consistency of identified QTLs with previous studies

Because the present experiment was conducted only under heat-stress conditions, the identified QTLs were compared with previously reported loci to assess their consistency across populations and experimental conditions. Several genomic regions overlapped or were located near previously reported QTLs for reproductive traits, providing independent support for the loci identified here.

The strongest consistency was observed for flower number. On chromosome 1, previously reported *qFPI1* and *qIN1* near 68 Mb [8] overlapped our flower-number QTL at 62.16–68.77 Mb. On chromosome 2, our QTL at 45.23–45.49 Mb co-localized with *nflw2.3* [21] and was also close to the heat-responsive *fln2.1_T3_2E* identified in an interspecific *S. lycopersicum* × *S. pimpinellifolium* population [20]. The repeated detection of this region across different genetic backgrounds provides strong support for its role in controlling flower number. Another notable region occurred on chromosome 6, where a QTL hub for fruit set and yield under heat stress was previously reported near 32.95 Mb [11]. In our study, QTLs for days to flowering, flower number, and fruit number were detected between 31.61 and 32.53 Mb. The convergence of multiple reproductive traits with a previously reported heat-responsive locus suggests that this region may contribute to reproductive performance under high temperature.

In addition, for pollen viability, *qPV11*, previously mapped near solcap_snp_sl_36066 at approximately 2.5 Mb on chromosome 11 under heat stress [8], was located near our QTL at 5.09 Mb. Notably, Nagcarlang was used as a heat-tolerant parent in both studies, suggesting that these loci may represent the same or closely linked genomic region. For days to flowering, the previously reported *flw3.3* [21] was located near our QTL at 58.37–59.83 Mb on chromosome 3, supporting the involvement of this region in flowering-time variation.

Overall, the alignment with previous studies provides independent support for several QTLs detected under heat stress, particularly the chromosome 2 region at approximately 44–45.5 Mb for flower number and the chromosome 6 region at approximately 31–33 Mb for multiple reproductive traits. These regions represent strong candidates for further validation and candidate-gene analysis.

## Discussion

### Key yield components under greenhouse and open-field conditions

The World Vegetable Center has conducted long-term breeding for heat-tolerant tomatoes adapted to tropical production systems, where open-field cultivation exposes plants to considerably greater environmental variation than controlled greenhouse conditions. Reproductive traits, particularly pollen viability and fruit set, have traditionally been used as indicators of heat tolerance in tomato because male reproductive development is highly sensitive to elevated temperature. Much of the experimental evidence supporting pollen viability as a selection criterion, however, has been generated under greenhouse, growth-chamber, or other controlled-temperature conditions. For example, heat stress under controlled conditions substantially reduces pollen viability and fruit set in heat-sensitive genotypes, whereas tolerant genotypes maintain higher pollen viability [8, 20]. A major QTL for pollen viability was also identified and validated under continuous mild heat, explaining approximately 36% of the phenotypic variation [8].

Our results suggest that the relative importance of these reproductive traits changes under open-field conditions. Among parental lines previously selected for heat tolerance under greenhouse conditions, only two maintained high fruit set in the field, indicating that reproductive performance identified under controlled conditions does not necessarily translate directly to the target production environment. Under field conditions, plants experience fluctuations in temperature together with other environmental factors that can affect successive stages of reproduction, including flowering, pollination, fertilization, and flower and fruit retention. Indeed, previous comparisons between controlled screening and field performance have highlighted the importance of validating physiological heat-tolerance indicators under field conditions.

This distinction was evident in the relationships among reproductive traits. Pollen viability showed moderate heritability (0.56) but was only weakly correlated with fruit set (*r* = 0.22), suggesting that pollen thermotolerance alone did not adequately predict realized reproductive success under open-field heat stress. In contrast, fruit number showed moderate heritability (0.48) and was strongly correlated with fruit set (*r* = 0.86), whereas fruit set itself exhibited negligible heritability (<0.01). These contrasting patterns suggest that fruit set is highly sensitive to environmental variation, while fruit number integrates the successful progression through several reproductive stages, from flowering and fertilization to early fruit retention.

Our results therefore do not diminish the importance of pollen viability as an indicator of reproductive thermotolerance; rather, they suggest that the most informative selection criterion may depend on the target production environment. Pollen viability provides a relatively heritable and physiologically informative trait for controlled screening, whereas fruit number may better represent realized reproductive performance under variable open-field conditions. Flower number could provide an additional complementary criterion because it showed the highest heritability among the reproductive traits (0.64). Thus, combining controlled-environment screening for specific physiological components such as pollen thermotolerance with field-based selection for fruit number and other reproductive outcomes may provide a more effective strategy for breeding heat-tolerant tomatoes adapted to tropical open-field production.

### Candidate genes underlying reproductive performance under heat stress

Because linkage disequilibrium in tomato can extend over relatively long and heterogeneous genomic regions, causal genes cannot be confidently assigned from GWAS signals alone. We therefore distinguished direct candidates, in which significant SNPs intersected annotated genes, from positional candidates located within 5 Mb of the corresponding QTL. Among these genes, two biological processes were particularly prominent: regulation of reproductive development and maintenance of cellular and protein homeostasis under stress.

### Regulation of floral and reproductive development

Several positional candidates have established functions in tomato reproductive development. *ARGONAUTE7* (*SlAGO7*; *Solyc01g010970*) is located approximately 0.85 Mb from the chromosome 1 SNP associated with days to flowering. *SlAGO7* regulates the trans-acting siRNA pathway, and its overexpression alters tomato inflorescence architecture and increases fruit yield [40]. Similarly, *EXCESSIVE NUMBER OF FLORAL ORGANS* (*ENO*; *Solyc03g117230*) lies within the chromosome 3 QTL region and regulates floral meristem size and floral organ number through the CLAVATA–WUSCHEL regulatory network [41]. On chromosome 2, *SlWUSCHEL* (*SlWUS*; *Solyc02g083950*) is located close to the flower-number QTL at approximately 45 Mb. Regulatory polymorphisms near *SlWUS* underlie the classical *locule number* (*lc*) locus and affect floral meristem activity and locule number [42]. Together, these positional candidates suggest that variation in floral meristem and inflorescence development may contribute to reproductive performance under heat stress. This hypothesis is particularly relevant because high temperature can disrupt floral meristem termination in tomato through regulation of *SlWUS*, resulting in abnormal flower and fruit development [43].

### Intracellular trafficking and protein homeostasis

Several direct candidates point toward a complementary role for cellular homeostasis. A significant SNP within the chromosome 3 days-to-flowering QTL intersected *Solyc03g115810*, encoding the vacuolar trafficking protein MON1. MON1 forms a complex with CCZ1 that activates Rab7-family GTPases and regulates prevacuolar compartment-to-vacuole trafficking and vacuole biogenesis [44]. MON1/CCZ1 from the wild tomato *Solanum chilense* has also been shown to enhance salt-stress tolerance when expressed in *Arabidopsis*, supporting a broader role for this trafficking pathway in abiotic-stress responses [45]. Although its role in heat-stressed reproductive tissues remains unknown, the direct association of *Solyc03g115810* makes MON1 a strong candidate for further investigation.

Protein turnover was also implicated by direct candidates on chromosomes 11 and 6. Significant SNPs within the chromosome 11 days-to-flowering QTL intersected *Solyc11g062310*, encoding an ECM29-like proteasome-associated protein. ECM29 regulates the 26S proteasome and promotes its disassembly in response to oxidative stress [46], suggesting a potential role in maintaining protein homeostasis during stress. More notably, the chromosome 6 region at approximately 31.6–32.5 Mb was associated with days to flowering, flower number, and fruit number and contained significant SNPs within ubiquitin-related genes, including *Solyc06g051090* and *Solyc06g051230*. The latter encodes an RMA1H1-like E3 ubiquitin ligase. RMA1H1 regulates aquaporin abundance and trafficking through ubiquitination, and its overexpression enhances drought tolerance [47]. Although these functions have not been directly demonstrated under reproductive heat stress, the convergence of multiple reproductive traits and several protein-regulatory genes within the chromosome 6 QTL suggests that protein homeostasis may contribute to reproductive stability under high temperature.

Overall, the candidate genes point to two complementary mechanisms: regulation of floral development, represented by positional candidates *SlAGO7*, *ENO*, and *SlWUS*, and maintenance of cellular and protein homeostasis, supported by direct candidates including MON1, ECM29, and the chromosome 6 ubiquitin-related genes. Among these, the chromosome 6 QTL is particularly noteworthy because it was associated with multiple reproductive traits, overlapped a previously reported heat-responsive QTL region, and contained several direct candidate genes. Nevertheless, these genes remain candidates for functional validation rather than confirmed causal determinants of heat tolerance.

### Applicability and limitations

The MAGIC population used in this study combines allelic diversity from multiple parental lines with increased historical recombination, providing advantages over conventional biparental populations for dissecting complex reproductive traits. The reduced confounding between population structure and relatedness in MAGIC populations can also facilitate the detection of marker–trait associations [22]. Importantly, several QTLs identified in the present study co-localized with loci previously detected in independent tomato populations, including the chromosome 2 region at approximately 44–45 Mb associated with flower number and the chromosome 6 region at approximately 31–33 Mb associated with multiple reproductive traits. The recurrence of these regions across different genetic backgrounds provides additional support for their potential relevance to tomato breeding. Nevertheless, the transferability of individual markers should be validated in independent breeding populations before their routine application in marker-assisted selection.

Several limitations should be considered when interpreting these results. First, the population was evaluated under open-field heat-stress conditions without a corresponding non-stress control environment. Consequently, the detected associations represent genetic variation in reproductive performance under high-temperature field conditions but cannot unequivocally be attributed to heat-specific responses. The overlap of several QTLs with previously reported heat-responsive loci provides independent support for their relevance, but multi-environment experiments including both optimal- and high-temperature conditions will be necessary to quantify QTL × environment interactions and distinguish constitutive effects from heat-specific responses. Second, despite the increased recombination provided by the MAGIC design, several QTL intervals remain relatively broad because of the extended and heterogeneous linkage disequilibrium in tomato. The candidate genes identified here should therefore be regarded as hypotheses rather than causal genes. Fine mapping, haplotype analysis, and functional validation will be required to resolve these regions and establish causal relationships. Integration with transcriptomic analyses of contrasting genotypes under control and heat-stress conditions could further prioritize candidate genes and clarify the molecular mechanisms underlying reproductive thermotolerance.

## Conclusions

This study provides new insights into the genetic and phenotypic basis of reproductive performance in tomato under open-field heat stress. Evaluation of a genetically diverse MAGIC population revealed substantial differences among reproductive traits in their heritability and relationships with reproductive success. Although pollen viability has been widely used as an indicator of heat tolerance under controlled conditions, its relatively weak association with fruit set suggests that it may not fully predict reproductive performance under variable field environments. In contrast, fruit number showed moderate heritability and a strong association with fruit set, supporting its use as a practical selection criterion for open-field heat tolerance. GWAS identified multiple QTLs for reproductive traits, several of which overlapped previously reported loci. Particularly notable were regions on chromosomes 2 and 6, with the chromosome 6 QTL associated with multiple reproductive traits. Candidate-gene analysis further implicated floral developmental regulation, intracellular trafficking, and protein homeostasis as potential mechanisms underlying reproductive responses to high temperature. Collectively, these findings emphasize the importance of evaluating heat tolerance in target production environments and provide genomic regions and candidate genes for further validation. Multi-environment evaluation, fine mapping, and functional studies will be essential to translate these findings into robust molecular tools for breeding heat-tolerant tomatoes.

## List of abbreviations

AF6: AmphaFluid 6
DF: days to flowering
FLN: flower number
FRN: fruit number
FS: fruit set
G × E: genotype × environment
GEMMA: Genome-wide Efficient Mixed Model Association
GWAS: genome-wide association studies
MAGIC: Multi-parent advanced generation intercross
PCoA: Principal coordinate analysis
PV: pollen viability
QTL: quantitative trait locus
SNP: single nucleotide polymorphism
TYLCD: tomato yellow leaf curl disease

## Declarations

### Ethics approval and consent to participate

Not applicable

### Consent for publication

Not applicable

## Availability of data and materials

The sequencing data underlying this article are available in NCBI at http://www.ncbi.nlm.nih.gov/bioproject/1210015, and can be accessed with BioProject PRJNA1210015. All data generated or analysed during this study are included in this published article and its supplementary information files.

## Authors’ contributions

YPL and PH: conceptualization and funding acquisition; YPL, KH, and EH: formal analysis; YPL, YCH, EH, CYL, YSC, CYL, LHH, JYY, PH, and KH: investigation; YPL, YCH, EH, SMH, and JYY: data curation; YPL, EH and KH: writing - original draft; YPL, EH, AE, PH, and KH: writing - review & editing; YPL, EH, and CYH: visualization; YPL, AE, PH, SWK, and KH: supervision.

## Acknowledgments

We extend our heartfelt gratitude to Peter Hanson for his invaluable contributions during his nearly three decades of leadership in the tomato breeding team at the World Vegetable Center. His dedication and expertise were instrumental in the development of this MAGIC population. We thank T. Komatsuda for technical assistance. We also thank the long-term strategic donors to the World Vegetable Center: Taiwan, the United States, Australia, the United Kingdom, Germany, Thailand, South Korea, the Philippines, and Japan.

## Supplementary information files

Fig. S1 Correlations among reproductive traits in the tomato MAGIC population under open-field heat-stress conditions.

Fig. S2 Quantile–quantile (Q–Q) plots of genome-wide association analyses for reproductive traits in the tomato MAGIC population.

Table S1 Molecular markers used to characterize disease-resistance genes in the parental lines of the tomato MAGIC population.

Table S2 Mean phenotypic values of the tomato MAGIC population evaluated under open-field heat-stress conditions.

Table S3 Significant SNPs associated with reproductive traits and their directly associated genes identified by GWAS in the tomato MAGIC population.

